# Effects of enhanced insect feeding on the faecal microbiota and transcriptome of a family of captive common marmosets (*Callithrix jacchus*)

**DOI:** 10.1101/2021.08.05.455322

**Authors:** Yumiko Yamazaki, Shigeharu Moriya, Shinpei Kawarai, Hidetoshi Morita, Takefumi Kikusui, Atsushi Iriki

## Abstract

Common marmosets have been widely used in biomedical research for years. Nutritional control is an important factor in managing their health, and insect intake would be beneficial for that purpose because common marmosets frequently feed on insects in natural habitats. Here, we examined the effect of enhanced insect feeding on the gut by analysing the faecal microbiota and transcripts of captive marmosets. A family consisting of six marmosets was divided into two groups. During the seven-day intervention period, one group (the insect feeding group, or Group IF) was fed one cricket and one giant mealworm per marmoset per day, while the other (the control group, or Group C) was not fed these insects. RNA was extracted from faecal samples to evaluate the ecology and transcripts of the microbiota, which were then compared among time points before (Pre), immediately after (Post), and two weeks after intervention (After) by total RNA sequencing. The gut microbiota of marmosets showed *Firmicutes*, *Actinobacteria*, *Bacteroidetes*, and *Proteobacteria* as dominant phyla. Linear discriminant analysis showed differential characteristics of microbiota with and without insect feeding treatment. Further analysis of differentially expressed genes revealed increases and decreases in *Bacteroidetes* and *Firmicutes*, respectively, corresponding to the availability of insects under both Post and After conditions. Significant changes specific to insect feeding were also detected within the transcriptome, some of which were synchronized with the fluctuations in the microbiota, suggesting a functional correlation or interaction between the two. The rapid changes in the microbiota and transcripts may be deeply connected to the original microbiota community shaped by marmoset feeding ecology in the wild. The results were informative for identifying the physiological impact of insect feeding to produce a better food regimen and for detecting transcripts that are currently unidentifiable.

## Introduction

After successful confirmation of the germline transmission of transgenes [1], common marmosets (*Callithrix jacchus*) have been increasingly used in various medical and biological areas [2]. Breeding methods for captive marmosets have been well established [3, 4, 5], while some health problems, such as diarrhoea and wasting, have been observed in many laboratories [6]. Marmoset wasting syndrome (MWS, or wasting marmoset syndrome, WMS) is a well-known health problem endemic to captive marmoset colonies and has been documented for several decades [7, 8]. The syndrome consists of various symptoms, but diarrhoea, anorexia, and anaemia are frequently observed [9, 10]. Several causes have been suggested to explain the variable symptoms, and malnutrition is thought to be one of the important factors for the aetiology of MWS [6, 7].

In the wild, common marmosets are known to maintain highly exudativorous (i.e., highly dependent on tree exudates, such as gum) diets [11], but they also feed on a variety of food items, such as insects, fruits, and small animals [12, 13]. Among them, insects are important nutritional resources because they account for 30-70% of their diet [3]. They eat various insects, such as grasshoppers, crickets, cicadas, and cockroaches [12]. Guidelines [3, 4] recommend providing captive marmosets with complete commercial food, insects and produce (vegetables and fruits). Although insects seem to have important nutritional roles in their health, the unique impact of insects on the physiological functions of marmosets has not been clarified. In the present study, we aimed to determine the effect of insect feeding on captive marmosets by analysing the microbiota and transcripts extracted from faecal samples.

The microbiota of common marmosets has been documented in several captive groups. One study [14] that compared the microbiota of faecal samples from individuals with and without WMS revealed differences in the abundance of only anaerobic, not aerobic, bacteria. The numbers of lactobacilli were lower in the WMS group than in the non-WMS group, whereas those of *Bacteroides*-*Fusobacteria* and *Clostridia* were higher in the WMS group than in the non-WMS group. The group with a higher rate of chronic diarrhoea had a lower proportion of *Bifidobacterium* than the other group, but there was no significant difference between the groups in terms of the Shannon diversity (H’) index [15]. Because daily feeding regimens vary at each facility, the microbiota should vary accordingly. However, there are still insufficient data to characterize the gut microbiota distribution of captive common marmosets. In the present study, we aimed to obtain basic data on the microbiota of marmosets by a total RNA sequencing (total RNA-seq) analysis method [16].

The total RNA-seq approach can be used to simultaneously describe microbial ecology and the transcriptome. It has been widely used in studies of marine ecology [17, 18], soil microbes [19, 20] and animal gut microbiota [21, 22] to obtain information from all domains of microbial inhabitants, including eucaryotes, archaea, and bacteria, without a strong PCR bias. Additionally, this approach can describe the gene expression patterns among samples, similar to standard transcriptome analyses, by using short-read alignment tools [23, 24, 25]. Therefore, we employed total RNA-seq to describe the activities of the whole microbial community in marmoset guts under our experimental conditions.

In the present study, we evaluated the effects of enhanced insect feeding for seven days on the gut microbiota and transcriptome by comparing groups with and without insect feeding. The main advantage of analysing both microbiota and transcripts simultaneously is to understand functional characteristics that would be attributable to ecological changes in the marmoset gut microbiota. The weekly weight and daily faecal scores were recorded to monitor the general health status throughout the experimental period. RNA was extracted from faecal samples from different timepoints (before (Pre), immediately after (Post), and after (After) the experimental intervention), and DNA was then sequenced and annotated with a database for taxonomic identification. In human subjects, the microbiota was reported to dramatically modify microbiota diversity after a change in diet for five days [26]. Thus, insect feeding is thought to have a substantial impact on the physiological states of marmosets, who preferentially eat insects in wild habitats [12].

## Materials and Methods

### Subjects

Six healthy adult common marmosets (*Callithrix jacchus*) from a family consisting of one mother (9 y) and five offspring (one male and three females, aged 3-4 y) were used in this study. The mean weight was 460 g, with a range from 374 g to 499 g. The mother was obtained from a company (Clea, Tokyo, Japan), and the offspring were laboratory born and raised by their own parents. They were living in a cage (w 70 x d 70 x h 180 cm) vertically separated by a metal mesh plate to prevent fighting; thus, they were physically separated but visually, acoustically, and olfactorily accessible to each other. The cage was placed in a breeding room on a 12-hour light-dark cycle and maintained at 28° and 50% of the temperature and humidity, respectively. According to this housing condition, the marmosets were divided into two groups that differed in terms of the amount of insect intake per week, as described below. After the study finished, the animals were not sacrificed, as the study did not include examination of postmortem specimens.

### Diets

The marmosets were fed commercially available pelleted foods (CMS-1M, Clea, Tokyo, Japan; SPS, Oriental Yeast, Tokyo, Japan) daily ad libitum in the morning and vegetables and fruits in the afternoon, in addition to a variety of food items such as yogurt, boiled eggs, acacia gum, cottage cheese, and small dried sardines. Different probiotic preparations (*Bifidobacterium bifidum* (Biofermine), Biofermin Seiyaku, Hyogo, Japan; *Bifidobacterium animalis* subsp. *lactis* (LKM512), Meito, Tokyo, Japan; *Bifidobacterium longum* and *Bifidobacterium infantis* (LAC-B), Kowa, Aichi, Japan) were added to the meals or given orally (1/2 to 1 tablet per head) when the animals had softened faeces or diarrhoea. Until the beginning of the current study, frozen house crickets (*Acheta domestica*), which were defrosted at the time of feeding, were given to all animals in the colony once per week (usually on Wednesday).

### Materials

For the insect feeding treatment, we used two different species, a house cricket (Tsukiyono farm, Gunma, Japan) and a giant mealworm (*Zophobas atratus*, Sagaraya, Kumamoto, Japan), which were commercially available and were kept frozen when they were delivered to the laboratory. They were brought back to room temperature to thaw just before feeding. These species have been reported to have similar amounts of protein, while the giant mealworm is much fattier than the cricket with higher calories [27].

### Procedures

#### Experimental design

The family was divided into two experimental groups, each with three subjects. One group (Group IF, consisting of three offspring females) was fed one cricket and one giant mealworm per day for seven continuous days. The other group (Group C, consisting of the mother, one offspring female, and one offspring male) was fed one cricket per week in the middle of the week, which was the regular food regimen in our colony. These insects were fed manually by the caretakers to each subject during the daytime.

There were three points of faecal sampling in the study: Pre, Post, and After. Pre samples were collected just before the one-week insect feeding period in the experimental group. Post samples were collected the day after the end of insect feeding, and After samples were collected two weeks after the insect feeding treatment.

#### Sample preservation

To analyse the microbiota and the transcripts of the faeces from the marmosets, samples were collected from the clean stainless floor of the breeding cages within 30 minutes of defecation early in the morning (8:30-10:30 AM) when they usually frequently defecated. Three pieces of faeces were collected from each group, and the faeces were immediately put into 10 ml RNAlater (Thermo Fischer Scientific, Waltham, MA, US) and manually mixed well with a sterile spatula to homogenize them in the liquid. Using the same procedure, three tubes consisting of three faecal pieces in 10 ml RNAlater were prepared for each group at each sampling point. The tubes were allowed to stand for 24 hours at room temperature. Then, they were stored at −80°C in a refrigerator until cDNA construction.

#### Faecal and insect RNA extraction, sequencing, and taxonomic annotation

Faecal RNA was purified by using the RNeasy PowerMicrobiota kit (Qiagen, Hilden, Germany). The kit was operated with an automatic system, QIAcube (Qiagen, Hilden, Germany), according to the protocol (RNA_RNeasyPowerMicrobiota_StoolOrBiosolid_IRTwithDNAse_V1.qpf) provided by the manufacturer. The concentration of RNA was measured with a Qubit 2.0 Fluorometer (Thermo Fischer Scientific, Waltham, MA, USA). For library construction, 10 ng of obtained RNA was processed using the SMARTer Stranded RNA-seq kit for Illumina (Takara Bio Inc., Shiga, Japan) according to the manufacturer’s instructions. After the concentration of DNA was evaluated by qPCR using the KAPA Library Quantification kit (KAPA Biosystems, Wilmington, MA, USA), the libraries were loaded onto an Illumina MiSeq sequencer and then sequenced using MiSeq Reagent kits v2 500 cycles (Illumina, San Diego, CA, USA) to obtain 250 bp paired-end reads. The nucleotide sequence data reported are available in the DDBJ/EMBL/GenBank databases under the accession numbers DRA008965 and DRA008966 for the marmoset and insect microbiota, respectively.

We used a mapping-based total RNA-seq pipelines [16] to analyse both rRNA and mRNA profiles to identify the taxonomy of the microbiota and to search for their functions. The obtained raw paired-end reads were trimmed by using Trimmomatic-0.35 [28] with seed mismatch settings: palindrome clip: simple clip threshold = 5:30:7, minimum read length of 100 bp, head crop of 6 bp and a specification to remove SMARTer kit-specific adaptor sequences. Then, trimmed paired-end reads were directly mapped to the SSU rRNA database SILVA release 128 rep-set data with 99% identity by Bowtie2 [23] with local mode default condition as a “best-hit” analysis. The data were transformed to BAM format for expression analysis. Mapped reads were counted by using eXpress [29] to obtain counting data against the SSU rRNA sequence database. Count data were combined with taxonomy data provided from SILVA release 128 (taxonomy all, 99% identity, taxonomy_7_levels.txt) by R [30].

To analyse the metatranscriptome, paired-end reads were assembled by using the Trinity v2.4.0 program package [31] with paired-end mode default settings. Open reading frames (ORFs) and the encoded protein sequences were predicted using Transdecoder. LongORF script in the TransDecoder v.3.0.0 program package (https://transdecoder.github.io/). The ORF data (longest_orfs.cds) were used as the reference database for read mapping. Mapping was performed as described above for SSU rRNA analysis. Functional annotation of the identified ORFs was conducted with the Trinotate-3.0.1 program package [32] (https://trinotate.github.io/). The obtained functional annotations were combined with read count data by R.

The obtained read count data were normalized according to Love et al. [33] using the TCC package in R. Additionally, SSU rRNA reads or ORFs with less than 10 mapped reads in total from all samples in the original count data were excluded by R.

#### Data analysis

The general health condition of the subjects was evaluated by weight and faecal score. The weight was measured once during each period of the experiment (i.e., Pre, Post, and After) as a part of the weekly physical examination performed by our laboratory. The faecal scores were measured daily by visual inspection of the faecal shape based on three levels (partially adopted from [34]): 3 corresponded to normal faeces (solid, with little liquid), 2 corresponded to loose faeces (globules with liquid but still formed), and 1 corresponded to diarrhoea (mostly globules, a large amount of liquid, and partially muddy).

The relative abundance of the communities with normalized read counts was analysed and visualized at the phylum and genus levels according to the groups (C and IF) and timing of the sampling (Pre, Post, and After). To evaluate the diversity/similarity of the microbes in the faecal samples, the Shannon and Chao1 indices were calculated and statistically analysed by using the RStudio software environment (ver. 1.3.1093 [35]). To see the similarity relationships among each condition of the groups in microbiota and transcripts, dendrograms were created by the clustering function in RStudio with the unweighted pair group method with arithmetic average (UPGMA) method of agglomeration using the Bray-Curtis index. For visualization of the distance of similarity among the conditions in spaces, nonmetric multidimensional scaling (nMDS) was conducted using the Bray-Curtis index, and the differences were statistically analysed by permutational multivariate analysis of variance (PERMANOVA).

To visualize the phylogenetic relationships of the microbes depending on the experimental conditions, we used the LEfSe program package [36] to conduct linear discriminant analysis (LDA) and to make a cladogram by following the instructions published online (https://huttenhower.sph.harvard.edu/galaxy/). For LDA, the LEfSe program was run with the threshold of 2.0 and an alpha value of 0.05 for both ANOVA (Kruskal-Wallis) and the Wilcoxon test.

To evaluate the changes in gene expression of microbiota and transcripts caused by the insect feeding treatment, differentially expressed genes (DEGs) were analysed using a pipeline (“EEE-baySeq”, 37), with a false discovery rate of 5%. To obtain DEGs, the datasets with groups (C and IF) and conditions (Pre, Post, and After) were divided into six groups (G1, G2, G3, G4, G5, and G6). G1, G2, and G3 corresponded to the Pre, Post, and After conditions of Group C, respectively. G4, G5, and G6 corresponded to those of Group IF. Because G1, G2, G3, and G4 were the conditions in which no insect feeding was executed, while G5 and G6 included insect feeding with different timings, any difference observed in G5 and G6 was of most interest in this study. Each group was compared to “others”, which included all conditions except for that group (e.g., G1 vs. others). The analysis also included the comparison among all the categories to find any differences (named “G7”: G1 vs. G2 vs. G3 vs. G4 vs. G5 vs. G6). Thus, eight DEG patterns were obtained involving six categories (no DEG, DEG G1, DEG G2, DEG G3, DEG G4, DEG G5, DEG G6, DEG G7). Then, for the microbes and transcripts associated with with any of the DEG patterns, the direction (i.e., larger or smaller than those of other categories) was determined: G1 > others, G2 > others, G3>others, G4>others, G5>others, G6>others, G1 < others, G2 < others, G3 < others, G4 < others, G5 < others, and G6 < others. For G7, only DEGs where G5 or G6 ranked among the top comparisons (i.e., G5>G3>G4>G6>G1>G2, for example) were considered for further analysis (G5>anywhere, G5<anywhere, G6>anywhere, and G6<anywhere). To overview the changes among the conditions, microbes with DEGs were presented at the phylum and genus levels. To detect the possible relationships, the microbiota and transcripts with DEGs were combinatorily clustered in each DEG category by using the ComplexHeatmap package in RStudio.

## Results

### General health condition

The faecal scores of the group subjected to the insect feeding treatment (Group IF) decreased (i.e., increase in the frequency of loose stools) in the After condition, as shown by the bars in Fig 1, but this was not statistically significant (*F* (2, 54) = 1.29, *p* = 0.283). Although the weights of the two groups were significantly different by 2-way ANOVA (*F* (1, 12) = 11.78, *p* = 0.005), they were stable during the whole experimental period, as shown by the lack of significance both among the conditions (*F* (2, 12) = 0.073, *p* = 0.930) and between the interaction of the group and condition (*F* (2, 12) = 0.011, *p* = 0.999), as shown in the lines in Fig 1.

**Fig 1.**
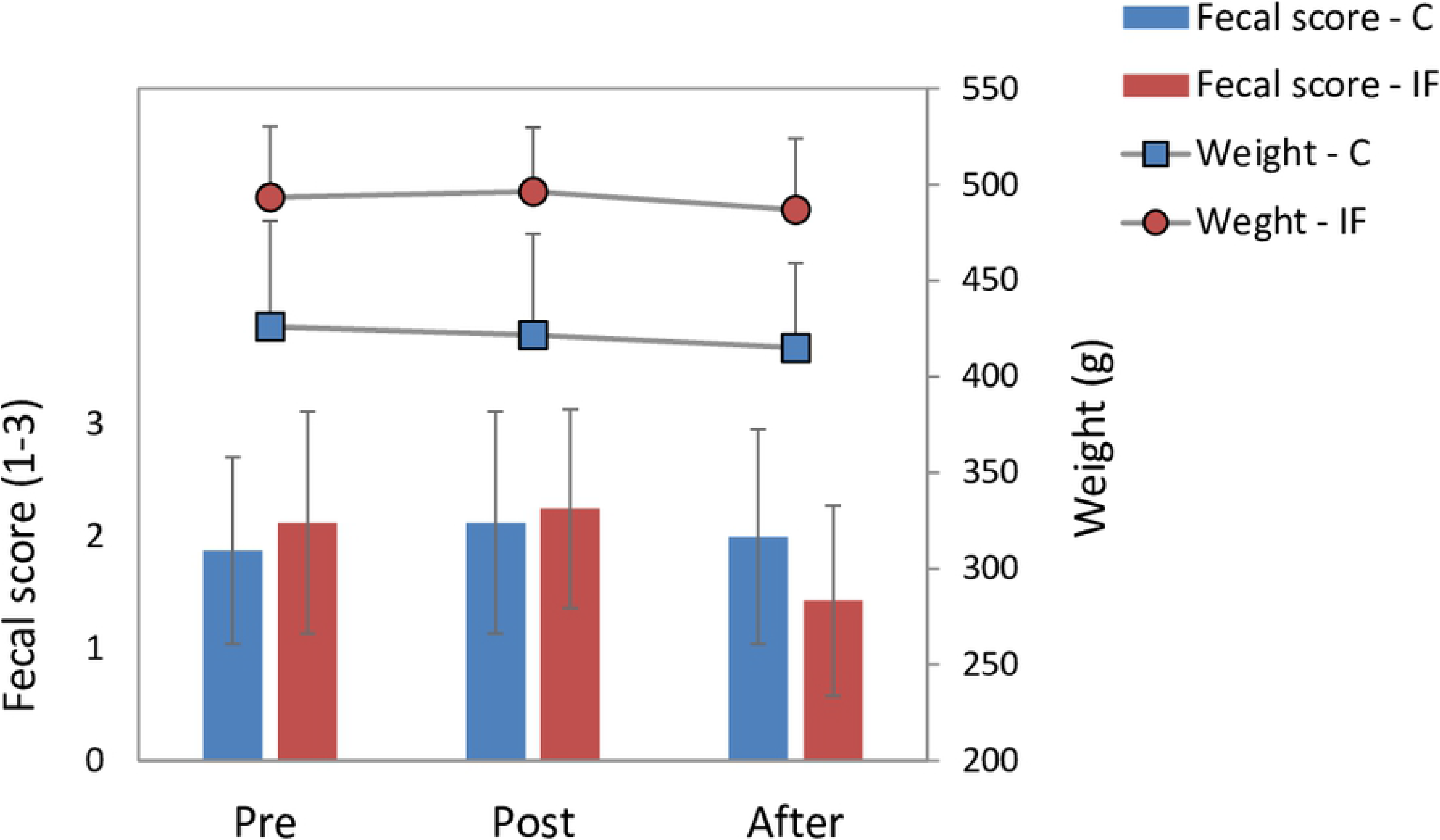
Faecal scores (bars) and weight (lines) during the experimental periods for Groups C (blue) and IF (red). Error bars show the standard deviation of the mean from three subjects. The faecal scores were recorded daily, and the weight was recorded individually once during each period.

#### Whole microbiota community of the marmoset gut

The total number of read counts normalized against the SSU rRNA sequences generated from the 18 faecal samples of the six common marmosets at the Pre, Post, and After conditions were 355,905.34, with an average of 19,772.52 counts per sample (see Supporting Information of S1 Table for the normalized count data of the annotated microbes). The difference in normalized read counts per sample between Groups C and IF was not significant (*t* (16) = −0.16, *p* = 0.88, mean ± standard deviation (SD) for Group C: 65.66 ± 2.63; Group IF: 66.03 ± 6.49).

Fig 2a shows the relative abundance of the microbiota at the phylum level for each group across the conditions. The phyla accounting for more than 0.5% of the total read count were listed, and those accounting for less than 0.5% and unable to be assigned to any phylum were categorized as “others”. *Firmicutes*, *Actinobacteria*, *Bacteroidetes*, and *Proteobacteria* were abundant in every sample of both groups. By observing the same data at the genus level, with more than 0.5% abundance, as shown in Fig 2b, *Veillonellaceae (*phylum *Firmicutes)* and *Bifidobacteriaceae* (phylum *Actinobacteria*) were dominant under all conditions.

**Fig 2.**
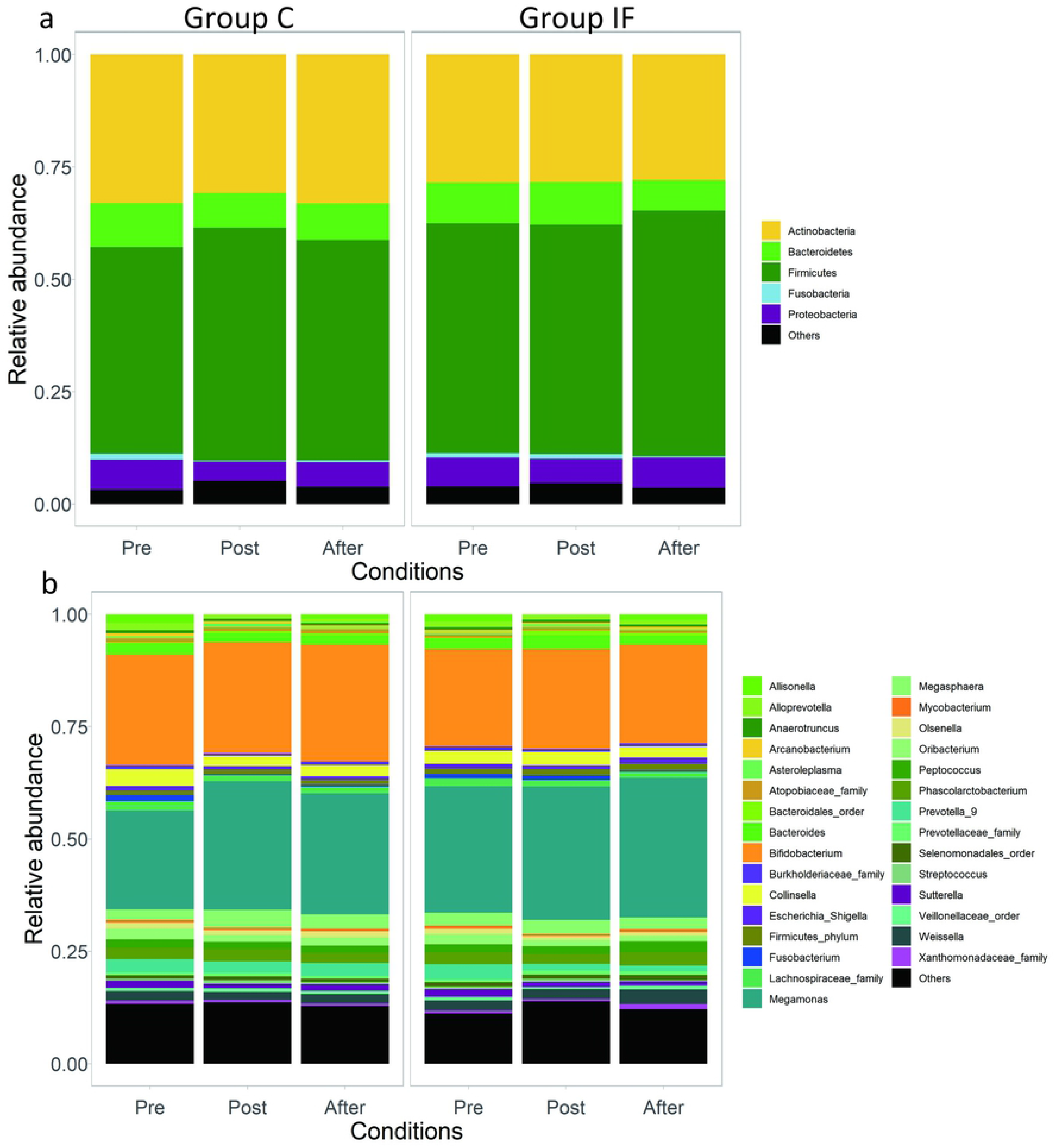
Relative abundance of microbes at the phylum (a) and genus (b) levels for Groups C (left) and IF (right) under the experimental conditions (Pre, Post, and After).

#### Microbial diversity under each condition

Shannon and Chao1 indices were used to determine the diversity of the microbial communities within the samples of each group, as shown in Fig 3. ANOVA of the Shannon indices after applying the general linear model (GLM) revealed a significant difference among the conditions (Pre, Post, After, F = 7.065, p = 0.008) but not between the groups (F = 0.292, p = 0.597). Similar results were obtained using the Chao1 index with the GLM (conditions: F=7.092, p = 0.007; groups: F=1.051, p = 0.323).

**Fig 3.**
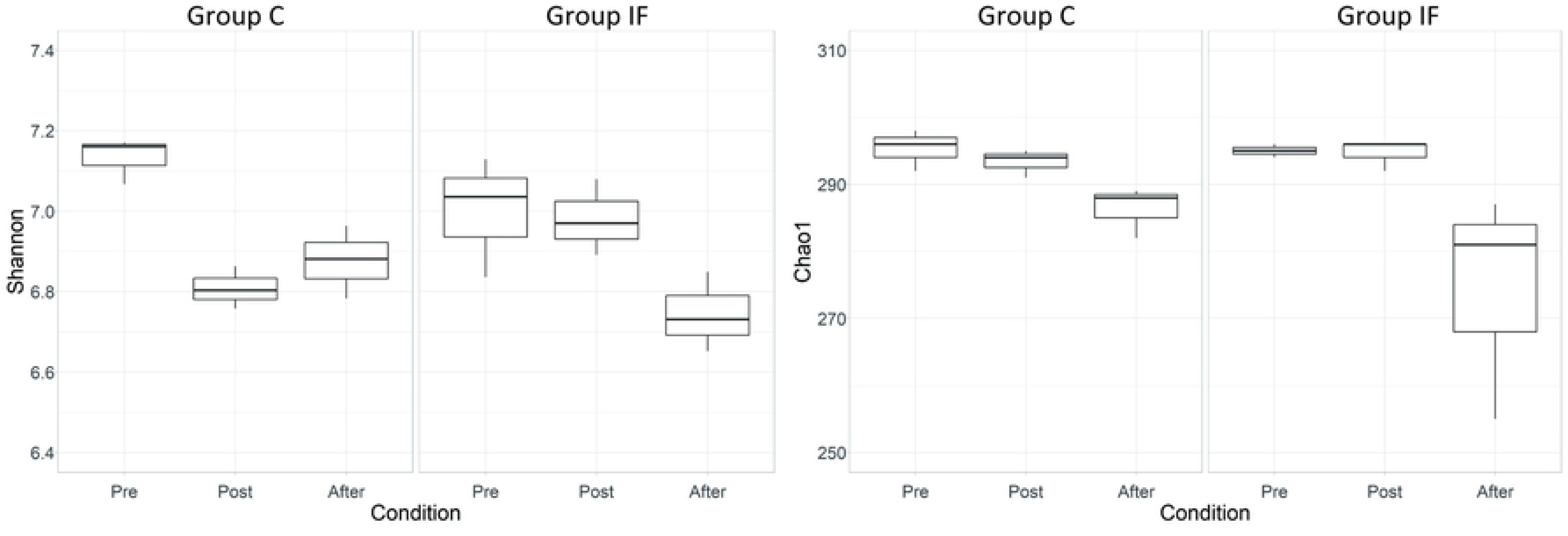
Shannon (left) and Chao1 (right) indices used to visualize the diversity within the samples of each group for the three conditions (left for Group C and right for Group IF in each panel, respectively).

#### Similarity of the microbial communities among the conditions

To determine the similarities of the microbial communities among the groups, the Bray Curtis index was calculated and used to generate the cluster dendrogram among the conditions of each group (Fig 4a), together with the differentially coloured squares and lines for individual conditions of two groups. For microbiota clustering, the Post conditions of Group IF (yellow solid line squares) were closely agglomerated.

**Fig 4.**
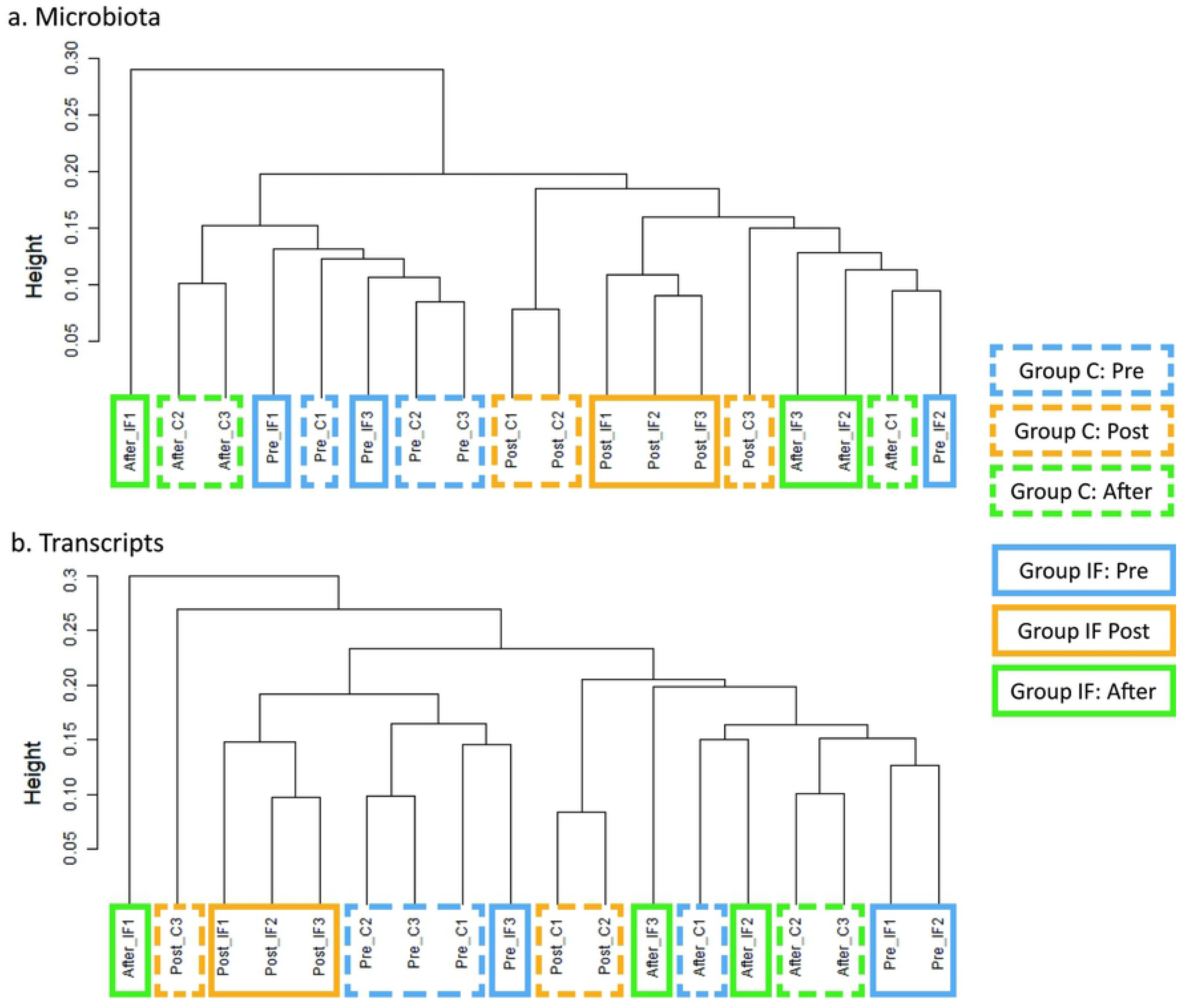
Cluster dendrograms for microbiota (a) and transcripts (b). Pre, Post, and After conditions are differentially coloured (sky-blue, yellow, and green), with dotted and solid lines for Groups C and IF, respectively.

To spatially visualize the microbiota similarities, nMDS was applied, and the data from each condition were shown in 2D space, as depicted in Fig 5a. To determine the effect of insect feeding, PERMANOVA was performed after dividing the individuals into three groups: (1) no insect feeding (three conditions for Group C and the Pre condition for Group IF), (2) Post of Group IF, and (3) After of Group IF, and significant differences were found among the three groups (F = 3.070, p = 0.005).

**Fig 5.**
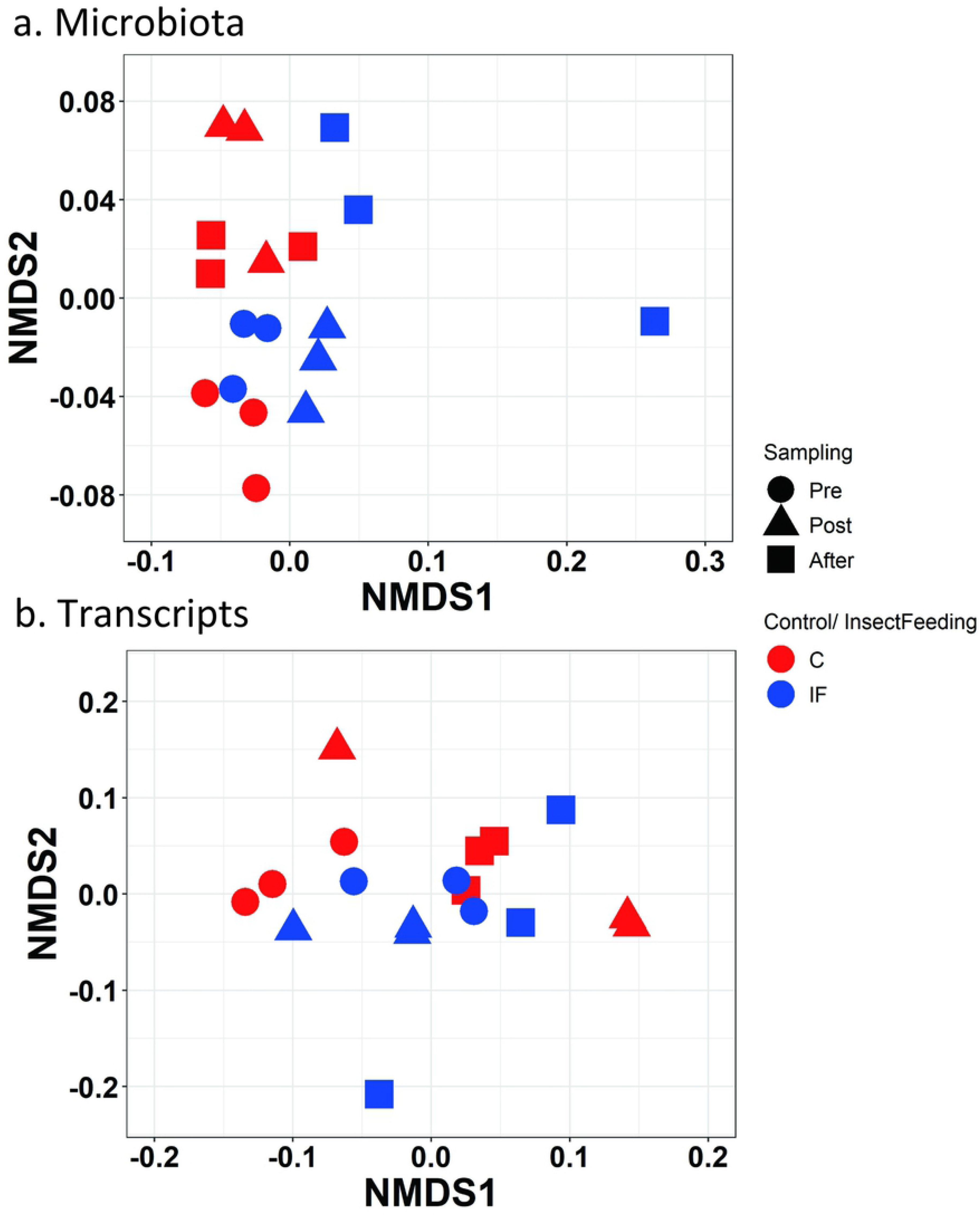
nMDS of the microbiota (a) and transcripts (b) data under the Pre (circle), Post (triangle), and After (square) conditions for Group C (red) and IF (blue).

#### Phylogenetic characterization of differentially abundant microbiota with and without insect feeding treatment

To characterize the specific microbes that emerged from the insect feeding treatment, we used the LEfSe analytical tool (see methods) on the normalized counts at multiple levels of taxonomy. The histogram in Fig 6a shows the LDA scores above the threshold (2.0) for the microbiota on various taxonomic levels ranked according to the effect size (alpha = 0.05) for the treatments with (IF: Post and After for Group IF) and without (NoIF: all conditions for Group C and Pre for Group IF) insect feeding. With IF, 5 taxonomic clades (three from the phylum *Proteobacteria* and two from *Bacteroidetes*) were differentially abundant. With NoIF, 8 taxonomic clades were detected by LDA (two from the phylum *Actinobacteria*, two from *Bacteroidetes*, and four from *Firmicutes*). Fig 6b shows the microbial abundance and taxonomic clades with differential characteristics for each treatment. The cladogram shows that abundant bacterial communities characteristic of IF were phylogenetically separatable from those characteristic of NoIF, as depicted by the red and green sectors without overlap.

**Fig 6.**
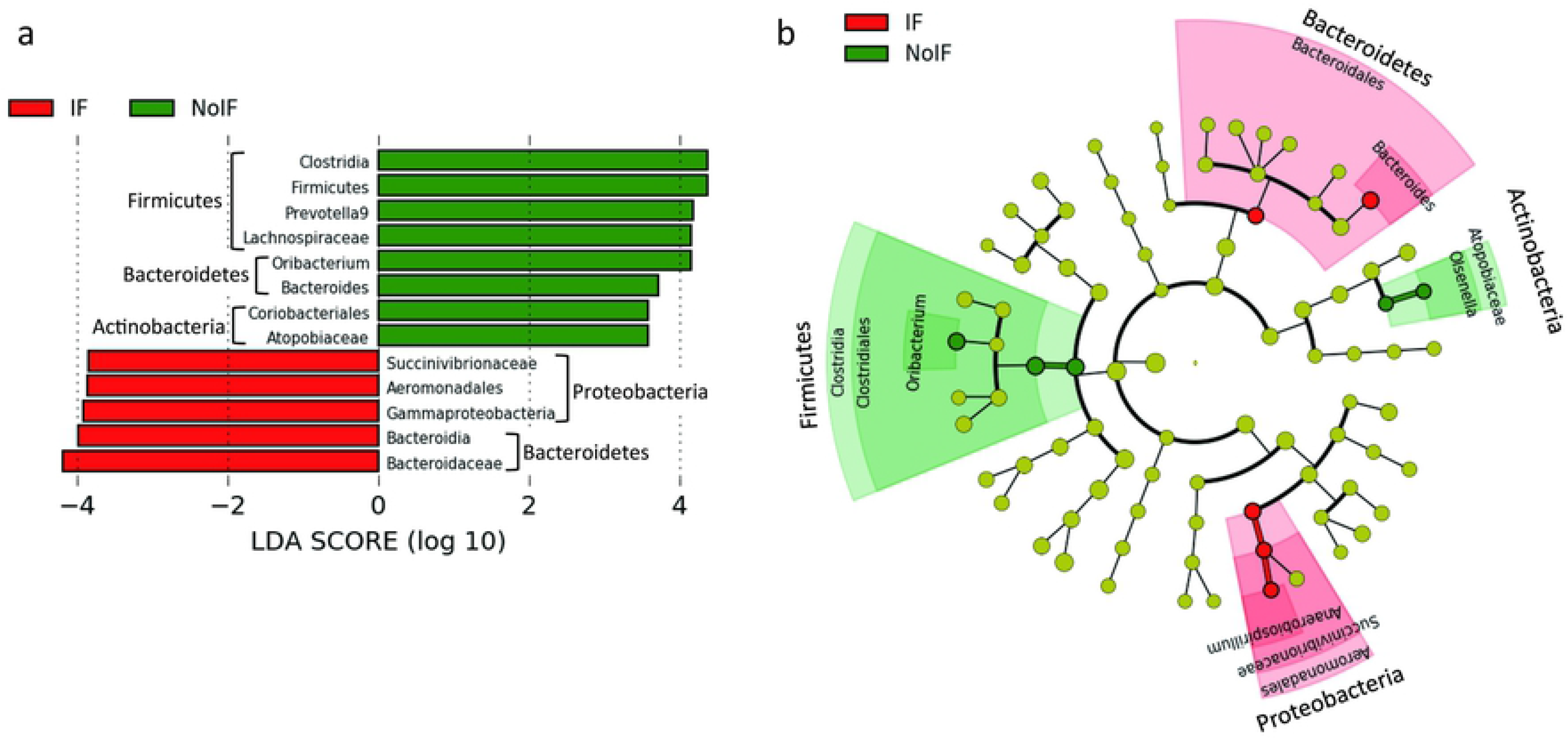
LEfSe characterization of the dominant microbial taxa according to the treatments with (IF: Post and After for Group IF) or without (NoIF: all conditions for Group C and Pre for Group IF) insect feeding. (a) LDA scores, above the threshold 2, of the microbial clades for each treatment, Insect feeding (red) and No insect feeding (green). (b) Cladogram based on the ranked list in (a) to visualize the relationships between the treatments and the phylogenetic relationships among the microbes. Phylogenetic clades are ordered from the centre of the circle, with narrower to broader taxonomic levels. Diameters of outer circles correspond to the relative abundance in the microbial community. Red and green points in the circles show the most abundant classes under the IF and NoIF treatments, respectively. Points in light green show the clades that are not significant.

#### Differentially expressed genes in microbiota among the conditions

The microbial communities fluctuated even without insect feeding treatment. Thus, to clarify the microbes specifically changed in Group IF depending on the conditions, the DEGs were analysed using the community data. Among 300 microbes annotated by the analysis, DEGs were confirmed for 99 of them. The relative distribution of DEG categories is presented in Fig 7a. Of 99 microbes with DEGs, a total of 21.21% were upregulated in terms of the experimental conditions (G5>others, G5>anywhere, G6>others, G6>anywhere), whereas a total of 20.20% were downregulated (G5<others, G6<others). No DEGs were found for the G3<>others, G5<anywhere, and G6<anywhere categories.

**Fig 7.**
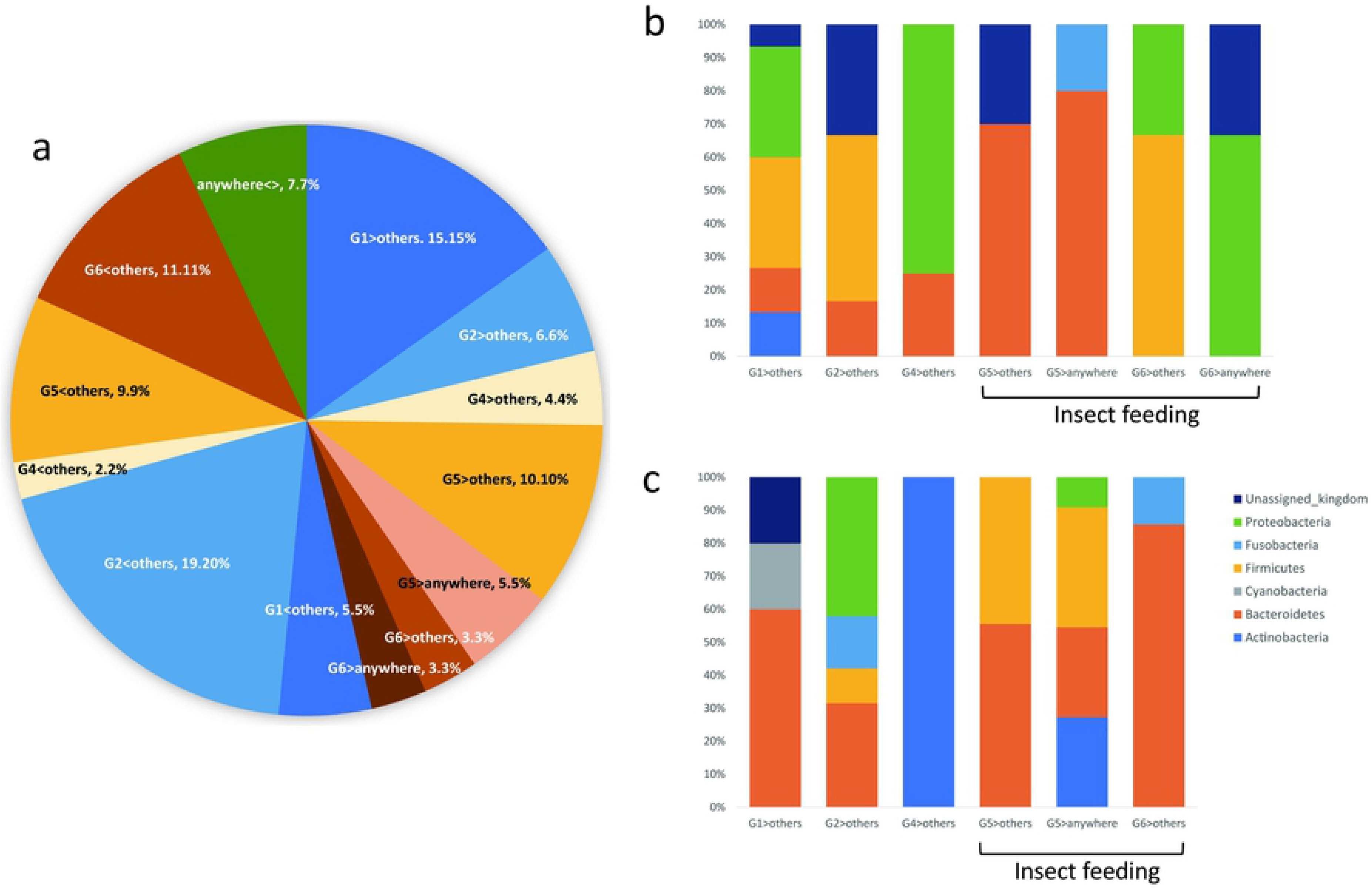
Distribution of DEG categories found in microbiota. (a) Relative distribution of the DEG categories. (b) Relative distribution of microbes in each DEG category at the phylum level, showing upregulated changes. Categories under the insect-feeding treatment were G5>others, G5>anywhere, G6>others, and G6>anywhere. (c) Relative distribution of microbes in each DEG category at the phylum level, showing downregulated changes. Categories under the insect-feeding treatment were G5>others, G5>anywhere, and G6>others.

Ninety-nine microbes with DEGs were further classified into phyla. Fig 7b and 7c show the microbes showing upregulated and downregulated changes with each DEG category, respectively. Among the upregulated phyla shown in Fig 7b, the distribution of *Bacteroidetes* was different under the Post condition of Group IF (i.e., G5>others and G5>anywhere). On the other hand, among the downregulated phyla in Fig 7c, Firmicutes showed different distributions in the same categories.

Fig 8 shows the normalized read counts of the microbes at the phylum level according to the DEG categories. The left two rows show upregulation, and the right row shows downregulation. In the case of the Post condition of Group IF (G5), *Bacteroidetes* appeared in both upregulated (Fig 8d, 8f) and downregulated (Fig 8k) categories, whereas *Firmicutes* only showed downregulated DEGs (Fig 8k). In the case of the After condition of Group IF (G6), *Actinobacteria* and *Bacteroidetes* only showed downregulated DEGs (Fig 8l), whereas *Firmicutes* and *Proteobacteria* showed both the upregulated (Fig 8e) and downregulated (Fig 8l) categories.

**Fig 8.**
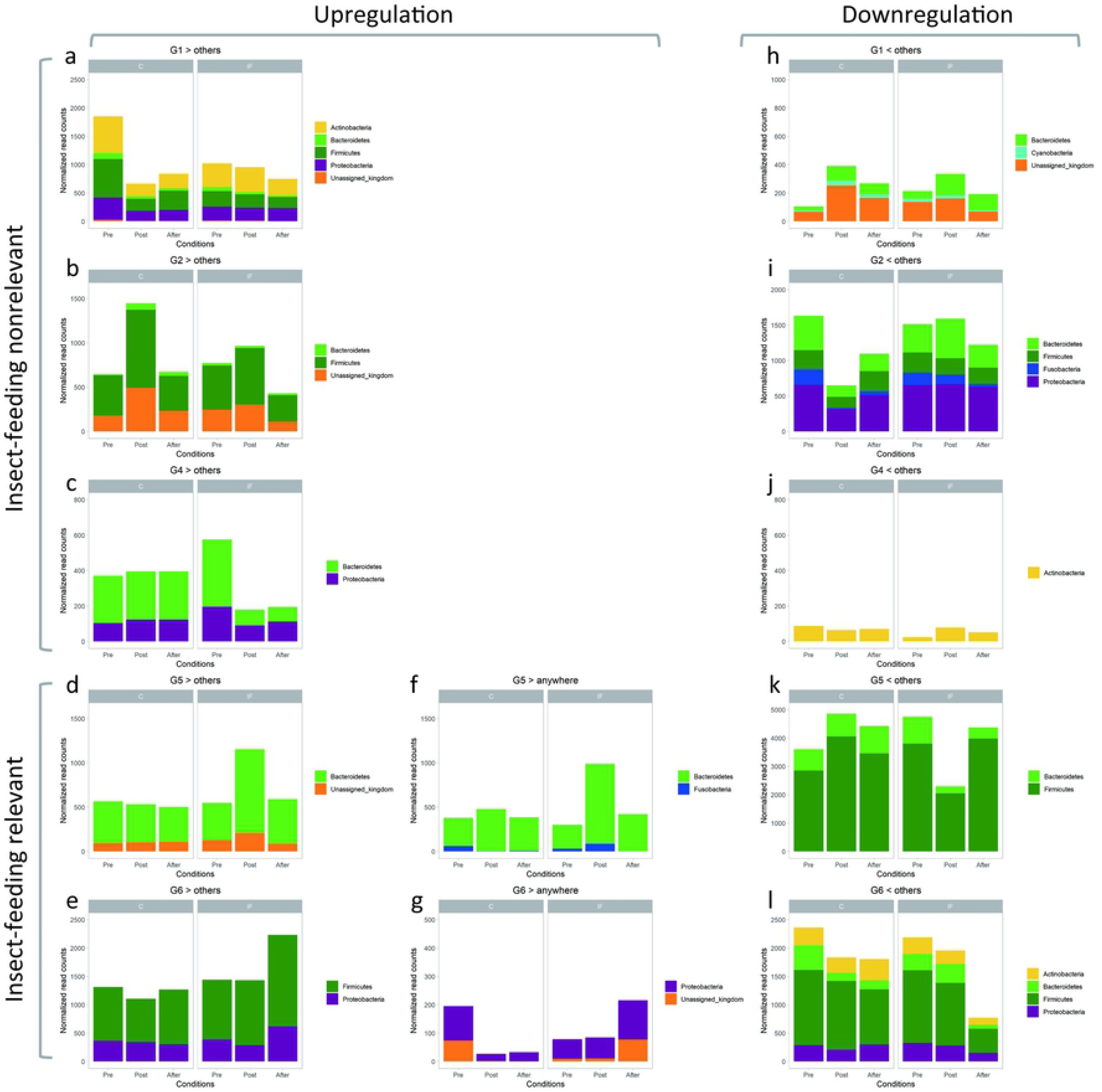
Normalized read counts of the microbes identified by the differentially expressed gene (DEG) analysis of faecal SSU rRNA at the phylum level. In each panel, the left and the right three bars show Pre, Post, and After conditions for Group C and Group IF, respectively. Black arrows in the panels indicate the groups which have DEGs compared with the other groups. (a-g) Upregulated DEGs, with insect-feeding nonrelevant (a, b, c) and insect-feeding relevant (d, e, f, g) conditions. (h-l) Downregulated DEGs with insect-feeding nonrelevant (h, i, j) and insect-feeding relevant (k, l) conditions. No DEG was found under the After condition in Group C (G3). For f, and g, data for each microbiota were compared with those of the other groups separately. In other graphs, data for each microbiota were compared with the total of the other groups.

To examine the changes according to the conditions (Pre, Post, and After) separately in the groups, Fig 9 shows the heatmap of the microbes at the genus level (for microbes unable to be annotated at the genus level, upper taxonomies (i.e., order, family) were assigned) with DEGs. The blue squares indicate the results relevant to the insect feeding treatment (Post (G5) and After (G6) conditions). In the upregulated category of the Post condition (G5>others, G5>anywhere), the genera *Bacteroides*, *Parabacteroides*, *Prevotella9* (phylum *Bacteroidetes*) and *Fusobacterium* (*Fusobacteria*) were listed, whereas in the After condition (G6>others, G6>anywhere), the genera *Weissella* (*Firmicutes*) and *Escherichia-Shigella* (*Proteobacteria*) were listed as having DEGs. In the downregulated category of the Post condition (G5<others), the genera *Bacteroides* (*Bacteroidetes*), *Allisonella, Megamonas,* and *Weissella* (*Firmicutes)* were found to have DEGs. In the After condition (G6<others), the genera were *Collinsella, Olsenella* (*Actinobacteria*), *Alloprevotella, Parabacteroides* (*Bacteroidetes*), *Streptococcus, Erysipelotrichaceae UCG-004, Oribacterium* (*Firmicutes*), and *Sutterella* (*Proteobacteria*).

**Fig 9.**
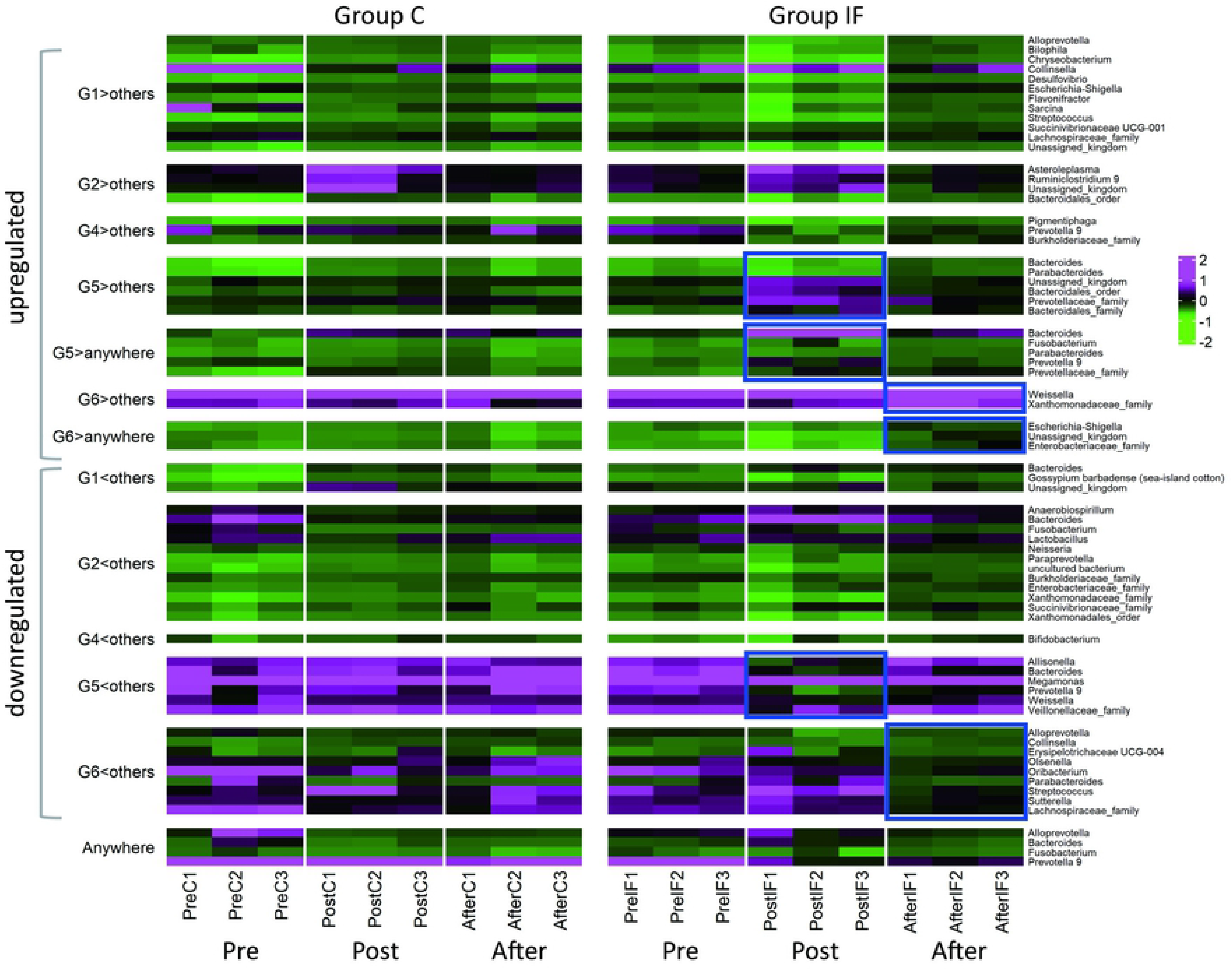
Heatmap of the SSU rRNA data (genus level) with DEGs. Groups C and IF are presented on left and right of the panel, respectively. For each group, the upper and lower parts present the data from upregulated and downregulated genera, respectively. G1: Pre condition for Group C; G2: Post condition for Group C; G4: Pre condition for Group IF, G5: Post condition for Group IF; G6: After condition for Group IF; AW: DEGs from any comparisons among the conditions. G3 is not presented because DEGs were not found under this condition (After condition of Group C). The heatmap with blue squares on the right indicates the changes in microbiota in accordance with the experimental treatments (i.e., G5 and G6 of Group IF).

LEfSe (Fig 6) revealed some genera with differential changes relevant to the insect feeding treatment, and thus we could identify the specific microbes of those genera by combining the results with the DEG results (i.e., selecting the DEG (G5 and G6) microbes in the genera found to be treatment-relevant by LEfSe). Fig 10 shows the normalized counts of such microbes from the genera *Bacteroides* (phylum *Bacteroidetes*), *Olsenella*, (phylum Actinobacteria) and *Oribacterium* (phylum *Firmicutes*). In *Bacteroides* (Fig 10a and 10b), there were microbes showing both upregulation and downregulation, but they were all relevant to the Post condition of Group IF (i.e., G5). On the other hand, the microbes in *Oribacterium* and *Olsenella* showed downregulation relevant to the After condition of Group IF (i.e., G6, Fig 10c and 10d, respectively).

**Fig 10.**
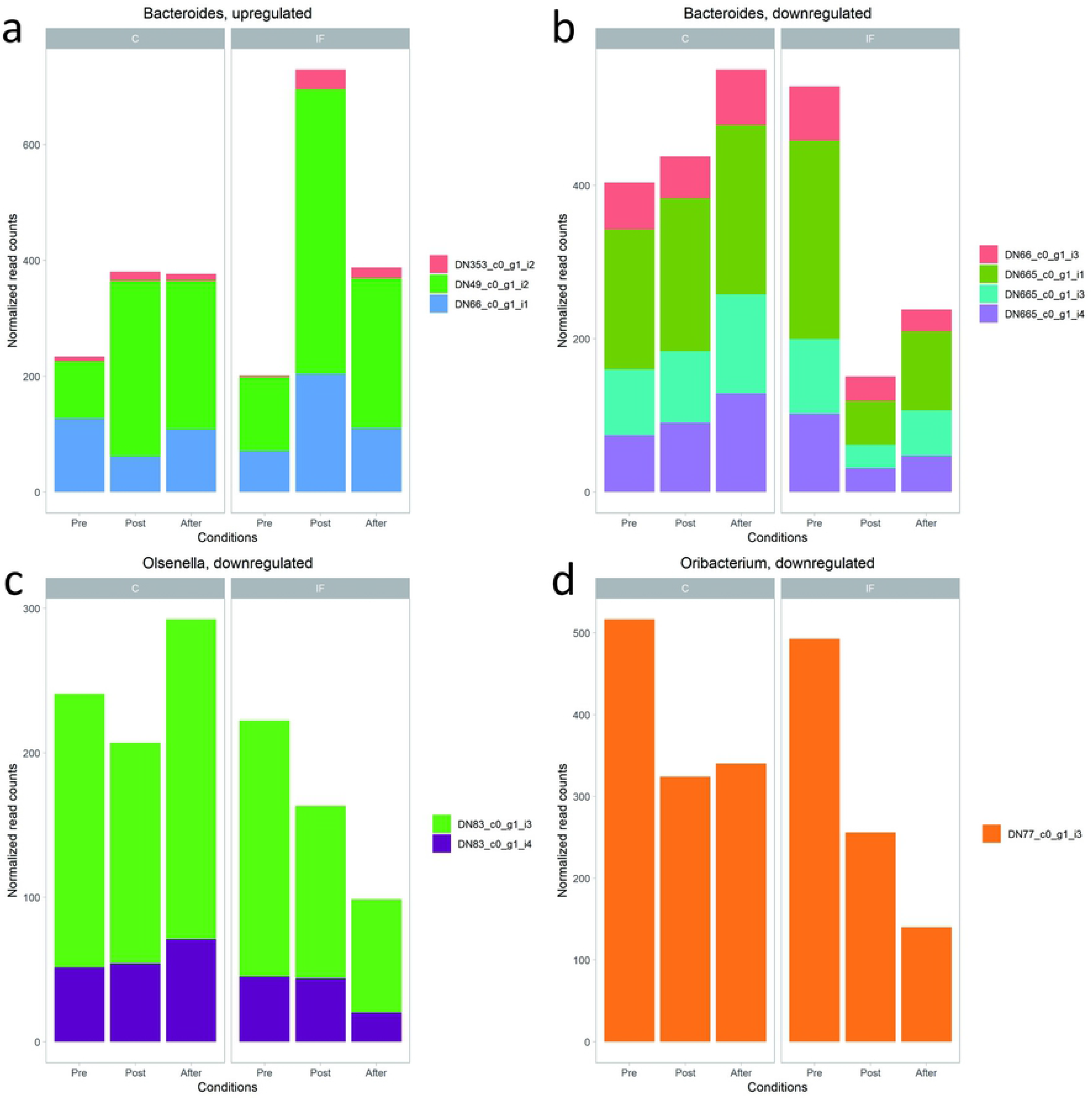
Results overlapped from the LEfSe and DEG analysis, showing the microbes specifically expressed just after the insect feeding treatment (a, b) and two weeks after the treatment (c, d), indicated by the black arrows. (a) Upregulation found under the Post condition in three microbes of the genus *Bacteroides*. (b) Downregulation found under the Post condition in four microbes of the genus *Bacteroides*. (c) Downregulation found under the After condition in two microbes of the genus *Olsenella*. (d) Downregulation found under the After condition in a microbe of the genus *Oribacterium*.

#### Transcriptomes of groups IF and C

Altogether, 407 different transcript IDs were classified by BLASTP (see Supporting information of S2 Table for normalized count data of transcripts). Among those, 72 were analysed with DEGs, among which 83% were classified as “hypothetical proteins” from various bacteria (Fig 11a). By classifying those proteins with an e-value above 1.0e+8, it was found that transcripts originating from *Bacteroidetes* and *Firmicutes* were abundant, as shown in Fig 11b and 11c. In the case of insect feeding-relevant conditions, both *Bacteroidetes* and unclassified bacteria were abundant in both upregulated (G5>others) and downregulated (G5<others) categories. *Firmicutes* characteristically increased in the downregulated category under the After condition (G6<others). *Proteobacteria* appeared only in the upregulated category under the After condition (G6>others).

**Fig 11.**
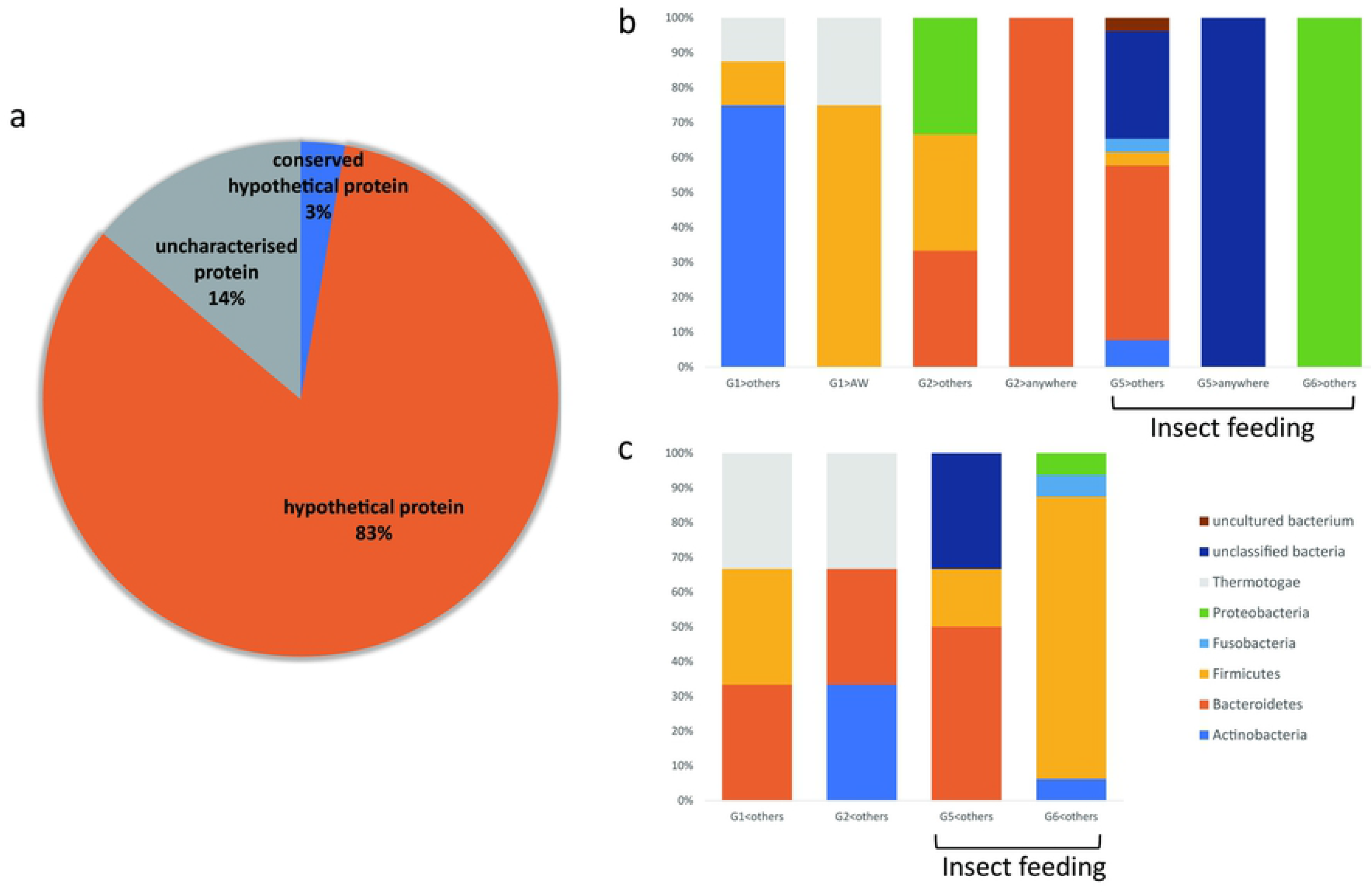
Relative distribution of DEG categories found in transcripts. (a) Relative distribution of transcript functions of DEGs. (b) Relative distribution of transcripts in each DEG category at the phylum level, showing upregulated changes. Categories under the insect-feeding treatments were G5>others, G5>anywhere, and G6>others. (c) Relative distribution of transcripts in each DEG category at the phylum level, showing downregulated changes. Categories under the insect-feeding treatment were G5>others and G6>others.

#### Relationship of the changes between the microbiota and transcriptome

The functional significance of transcripts was evaluated by analysing the similarity of changes between the microbiota and transcriptome and whether such changes closely interacted with each other according to the experimental conditions. Thus, we performed clustering analysis with Pearson’s product moment correlation coefficient among the SSU rRNA and transcriptome data with DEGs within 6 categories. Fig 12 shows the heatmap of the changes in the microbiota and transcriptome listed together in each DEG category, according to the experimental period (indicated on the bottom of the heatmap) in Groups C (left) and IF (right). The dendrogram located on the right of the heatmap shows the results of Group IF. Note that the microbiota and transcriptomes that behaved similarly were near each other in the heat map with a short clustering distance. The blue squares indicate the results relevant to the insect feeding treatment. In the case of the upregulated categories, microbes and transcripts tended to cluster separately. In the case of the downregulated categories, especially G6<others, they were more intermixed.

**Fig 12.**
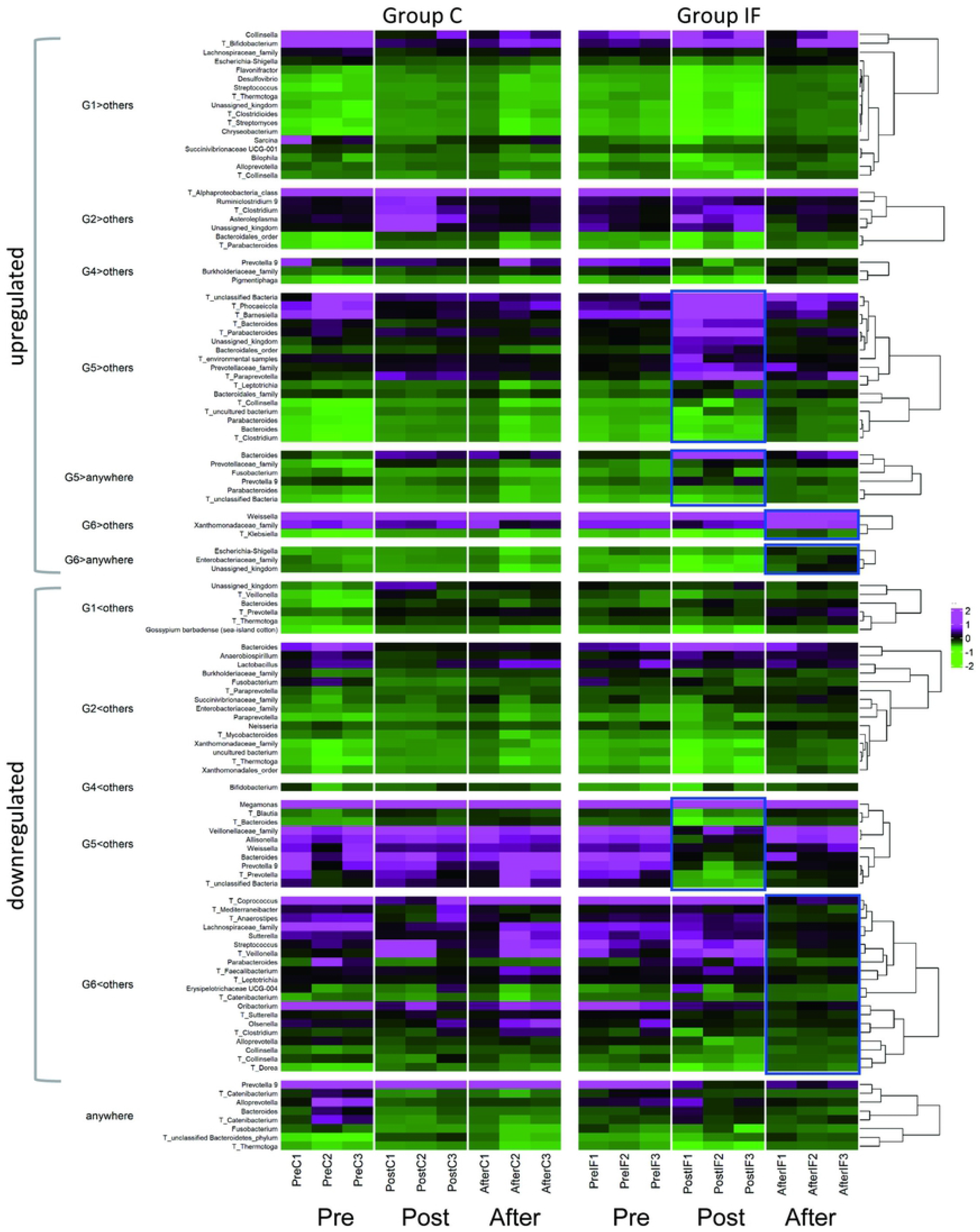
Heatmap with clustering among the SSU rRNA (genus level) and annotated transcripts with DEGs. Transcript data are presented with T with the genera name of the originating microbes. Increases and decreases are shown in magenta and green, respectively. Hierarchical clustering enabled items from the microbiota and transcriptome data to be listed together, where items with similar characteristics were listed close to each other. The data relevant to the experimental treatment (insect feeding) are emphasized by the blue squares. Upregulated and downregulated DEGs are located upper and lower panels of the figure, respectively.

#### Microbiota and transcripts of the insects fed to the marmosets

To determine whether the significant changes in the abundance of the microbial community in the samples of Group IF after insect feeding were attributable to the insects themselves fed to the marmosets, we analysed the microbiota of crickets and mealworms from the same lot as those fed to the subjects, using the same protocol described in the methods (see Supporting information of S3 and S4 Tables for normalized count data of the microbes and transcripts, respectively). As shown in Fig 13, abundant microbial phyla accounting for more than 0.5% of the total reads were *Opisthokonta* (eucaryotes), *Archaeplastida* (eucaryotes), and *Firmicutes* for the crickets and *Opisthokonta*, *Archaeplastida*, *Firmicutes*, *Proteobacteria*, and *Cyanobacteria* for the meal worms. The relative abundance of the kingdom *Bacteria* was 1.09 and 7.26% of the total reads from the crickets and worms, respectively, while it was 96.38% of the total reads from the faecal samples of marmosets. All the remaining microbes were in the kingdom *Eukaryota* (98.91% and 92.04% for crickets and mealworms, respectively), which was 0.28% in the case of the marmoset sample.

**Fig 13.**
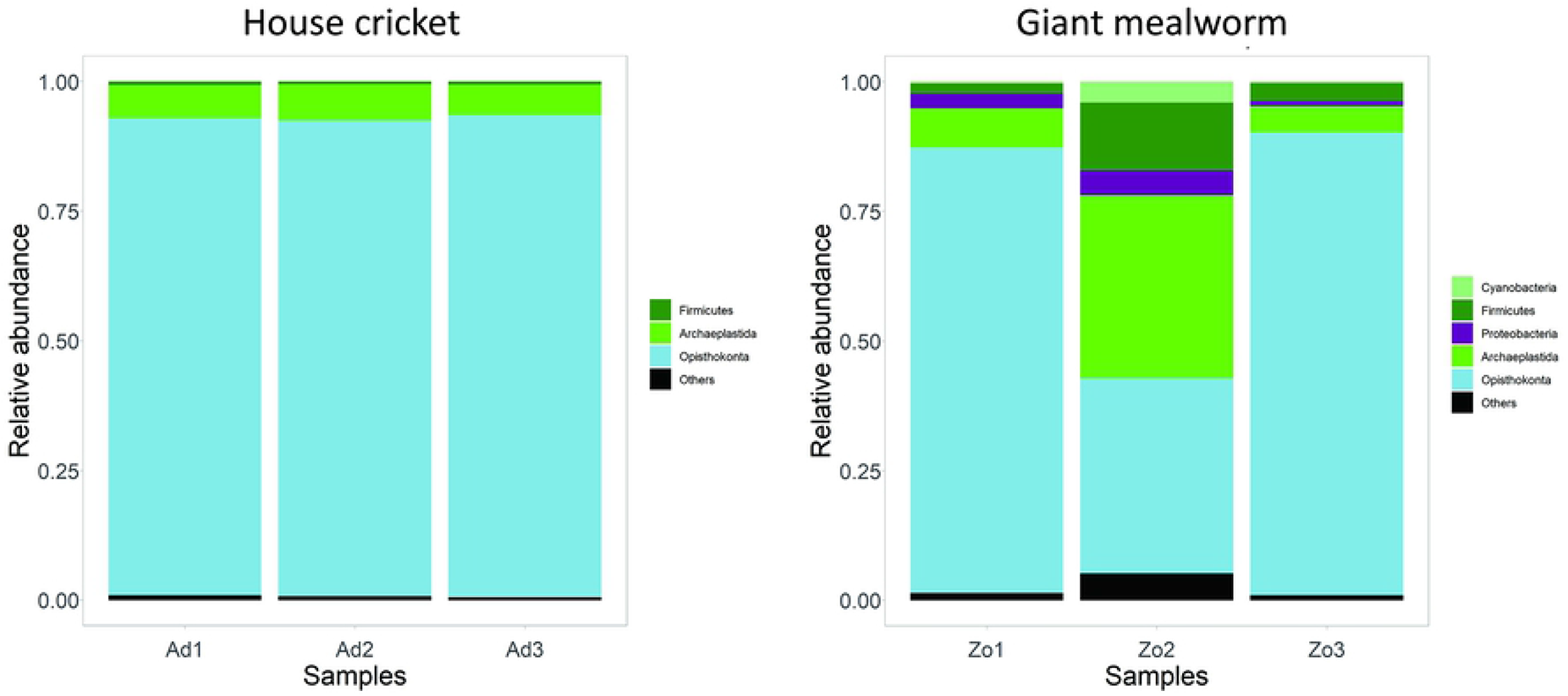
Relative abundance of microbes at the phylum level for house cricket (left) and giant meal worm (right).

## Discussion

The present study described the characteristics of the gut microbiota of captive common marmosets by using the total RNA sequencing method, with *Firmicutes*, *Actinobacteria*, *Bacteroidetes*, and *Proteobacteria* as dominant phyla. Then, we showed that enhanced insect intake for only one week modified the microbiota population in the gut, which interacted with the transcripts simultaneously extracted from the faecal samples. Changes observed in the microbiota were not attributable to the insects themselves. More specifically, microbes in the phyla *Bacteroidetes* and *Firmicutes* showed corresponding changes in their abundance under the insect feeding treatment at the different sampling points. *Bacteroidetes* showed both an increase and decrease upon just finishing the treatment (Post condition), followed by a decrease after two weeks (After condition), while *Firmicutes* showed a decrease at the Post condition, followed by both an increase and decrease at the After condition. These results corresponded well to the changes in the abundance of transcripts having the same homologous phyla of origin. Overall, the current study indicated that a partial change in the diet for seven days had an impact on the host marmosets’ microbiota and that insect feeding naturally observed in wild populations of common marmosets has special roles.

### Treatment-relevant changes observed in the microbes belonging to the phyla *Bacteroidetes* and *Firmicutes*

Treatment with insect feeding differentially affected the marmosets’ gut microbiota at different times. As indicated by LEfSe (Fig 6), sets of microbes in the phylum *Bacteroidetes* appeared in both categories, whereas those of the phyla *Proteobacteria* (insect feeding) and *Actinobacteria* and *Firmicutes* (no insect feeding) appeared in either of the categories. Additionally, DEG analysis with 6 patterns showed the exact changes according to the groups and conditions (Figs 8 and 9). Because the phylum Bacteroidetes showed both an increase and a decrease under Post conditions and a decrease under After conditions and the phylum *Firmicutes* showed a decrease under Post conditions and an increase and decrease under After conditions, one possible interpretation of these results would be that *Bacteroidetes* and *Firmicutes* had opposite responses to insect feeding. That is, microbes in those phyla may have shown rapid changes corresponding to the availability of insects. Adding one more sampling point during the insect feeding treatment would clarify the above possibility, which was not examined in the current study.

The results from the studies on other species treated with animal-concentrated diets would be comparable to those obtained in the current experiment. In a human study, after taking an animal-based diet for five days, an increase was found in species belonging to the genus *Bacteroides*, whereas a decrease was found in those of *Firmicutes,* but there were some species belonging to *Firmicutes* that showed increased in abundance after the treatment [26]. This study also detected an increase in the abundance of bacteria in the phylum *Proteobacteria*, including the species *Bilophila wadsworthia*, which is known to be stimulated by increased bile acid responsible for fat intake [38]. In our study, both an increase and a decrease in the abundance of *Proteobacteria* were observed under After, not Post conditions (Figs 8e, 8g, and 8l), suggesting a gradual change in the metabolism related to bile acid in the gut of the host. A previous study on dogs fed a raw meat diet for 14 days [39] observed 7 genera showing an increase after the treatment, and only the genus *Bacteroides* was in common with our current study findings. Chickens fed *Tenebrio molitor* larvae for 54 days showed an increase in *Firmicutes* and a decrease in *Bacteroidetes* at the phylum level [40], which corresponded to some of our results (Figs 8d, 8f, and 8e).

### General impact of insect feeding on the gut

The abundances at the phylum and genus levels, together with two indices of α diversity (Shannon and Chao1 indices), did not differ from each other between Groups C and IF. At the macroscopic level, insect feeding for seven days did not affect the general community of the intestinal microbiota, which was in common with the results of human adults treated with 25 g cricket powder per day for 14 days [41]. However, after focusing on the pattern of the changes corresponding to the experimental treatment by calculating the β diversity, there were significant differences between the conditions with and without insect feeding. The same patterns, no difference in α diversity but a significant difference in β diversity, were observed in a study on human subjects treated for 5 days with an animal-based diet [26]. A study examining the effects of an animal-based diet on the microbiota in dogs reported that faeces became firm and that the Shannon H’ index increased after raw beef was added to commercial food for 14 days [39]. The reason for the lack of a significant change in α diversity indices (Shannon and Chao1) after insect feeding in the current study would partially be attributable to unexpected fluctuations observed in the samples of Group C. Thus, evaluation of the stability of the microbial community would be necessary to compare the effects of short-term intervention (seven days) by taking additional samples in the Pre period, for example.

For microbiota, the samples just after the insect feeding treatment were clearly distinguishable from other samples, as shown in Fig 4a. By examining the distances among the samples, conditions with insect feeding (Post and After conditions of Group IF) were closely grouped and more distance from the other samples. Additionally, as the results of the PERMANOVA indicated, there were also differences between the Post and After conditions of Group IF, suggesting that changes caused by the insect feeding treatment had impacts on microbiota, which were long lasting for the bacterial community even after the treatment ended. On the other hand, PERMANOVA did not detect a significant difference in the transcript data between the conditions with and without insect feeding, although the samples under the Post conditions were closely clustered in Fig 4b, and the plots were separately located in Fig 5b. The reason for this difference between the microbiota and transcriptome was not identified by the current results, so some possibilities, such as length of the treatment and quantity of the fed insects, need to be examined to determine the changes in the transcripts.

### Limitations and future perspectives of the study

Our analysis using total RNA-seq, which could concurrently detect the dynamics of the microbiota and transcripts, was effective in searching for functional genes that are currently unidentifiable after establishing a metagenomic database of the transcriptome with a full-length cDNA library. However, the functional significance of the transcriptomes could only be inferred by the microbiota that showed similar changes across the experimental conditions. In the next step, we need to capture a wider view that would integrate the changes in microbiota, transcripts, and host responses to understand the effect of feeding insects, considering that the subjects have a long history of foraging.

In the present study, we used frozen crickets and giant mealworms instead of live mealworms. The results might have been different from those obtained in the study using live insects. The use of live insects as a feeding regimen is also recommended for marmosets in terms of enrichment purposes [4]. Nevertheless, the present study showed that feeding common marmoset insects changes their physiological status by balancing the microbiota to modulate metabolites. Consideration must be given to how much and what types of insects we should feed captive marmosets because there is a risk of overeating in the breeding cages but not in bushes in the wild. For example, some insect larvae are rich in fat and lack calcium, and it is therefore recommended to feed the insects a high calcium diet before feeding the insects to marmosets [3]. Insects with a high phosphorous-calcium rate should not be provided in abundance to prevent the malabsorption of calcium [4]. Thus, further studies are clearly needed to determine the long-term effect of insect feeding in captive animals and what types of insects are most beneficial for their health while simultaneously monitoring the changes in the faecal microbiota and transcriptome.

## Conclusions

The present study showed that adding insects to the regular food regimen for seven days could have a distinct effect on the microbiota and transcripts of captive common marmosets. The total RNA-seq method was used to analyse the microbiota and transcripts simultaneously, and the correlational analysis suggested that they did interact with each other. Thus, enhanced insect feeding could activate the physiological dynamics that have been evolutionarily developed in this species in wild habitats. The obtained results help us to understand the interaction between the host and the microbiota via food sources and suggest that the feeding ecology in wild habitats is an important key to developing food regimens appropriate for the microbiota of common marmosets.

## List of abbreviations

ANOVA: analysis of variance
Group C: control group.
Group IF: insect feeding group
LDA: linear discriminant analysis
LEfSe: linear discriminant analysis effect size
MWS: marmoset wasting syndrome
ORF: open reading frame
PERMANOVA: permutational multivariate analysis of variance
SSU rRNA: small subunit ribosomal ribonucleic acid
total RNA-seq: total RNA sequencing
WMS: wasting marmoset syndrome

## Declarations

### Ethics approval and consent to participate

This study was approved by the Animal Experiment Committees at the RIKEN Brain Science Institute (H27-2-203) and was conducted in accordance with the Guidelines for Conducting Animal Experiments of the RIKEN Brain Science Institute. Consent to participate was not applicable to the study.

## Consent for publication

N/A

### Availability of data and materials

The datasets supporting the conclusions of this article are included within the article and its additional files.

### Competing interests

Authors declare no competing financial interests.

### Funding

Grants of Brain Mapping by Integrated Neurotechnologies for Disease Studies (Brain/MINDS) at RIKEN from Japan Agency for Medical Research and Development (JP18dm0207001) and the Science Research Promotion Fund for Private University at Azabu University from the Ministry of Education, Culture, Sports, Science, and Technology of Japan supported the purchase of the materials for RNA extraction and preservation of faecal samples and the costs of sequencing.

### Authors’ contributions

Y.Y., S.M., and S.K. designed and executed the experiments, analysed the data, wrote the main manuscript text, and prepared the Figs. S.M. developed and applied the total RNA-seq pipeline for analysing microbiota and transcriptome data. S.K. supervised the health status of the subjects; H. M, T. K, and A.I. coordinated manuscript writing. All authors reviewed the manuscript.

## Acknowledgements

N/A

## Supporting information

**S1 Table. Normalized counts of microbes annotated by QIIME 2 vsearch for Groups C and IF.**

**S2 Table. Normalized counts of transcripts annotated by blastp for Groups C and IF.**

**S3 Table. Normalized counts of the microbes of the insects (house crickets and giant meal worm).**

**S4 Table. Normalized counts of the transcripts of the insects.**

## Notes

### Competing Interest Statement

The authors have declared no competing interest.

